# Dental biofilms contain DNase I-resistant Z-DNA and G-quadruplexes

**DOI:** 10.1101/2024.10.13.618059

**Authors:** Dominique C. S. Evans, Mathilde F. Kristensen, Lorena G. Palmén, Inge Knap, Manish K. Tiwari, Sebastian Schlafer, Rikke L. Meyer

## Abstract

eDNA is a major component of the extracellular matrix of bacterial biofilms, and recent studies have shown that biofilms from many pathogens contain both Z-DNA and G-quadruplex (G4) structures in addition to the canonical B-DNA double helix. These secondary DNA structures provide new emergent functions to the biofilm, most notably by making biofilms more resilient by protecting them from nucleases. In dental biofilms, it is largely unknown what conformation eDNA adopts, and the aim of this study was to determine if dental biofilms contain non-canonical secondary DNA structures.

In this study, we visualised B-DNA, G4, and Z-DNA in dental biofilms from 10 healthy subjects and from 10 caries-active subjects using fluorescence immunolabeling and confocal microscopy. eDNA formed large structures surrounding clusters of microorganisms that contained B-DNA, G4, and Z-DNA in the majority of the samples. We also identified microorganisms outside of these large eDNA structures that contained smaller G4 and Z-DNA structures associated to cell surfaces. G4 and Z-DNA are resistant to degradation by the commonly used mammalian DNase I. We verified this resistance in dental biofilms, and we suggest that these structures present a novel target for improved enzyme formulations for controlling oral biofilms and, more broadly, any biofilm that contains G4 and Z-DNA.

## Introduction

Dental biofilms are complex and diverse microbial communities encased in a protective extracellular matrix comprised of polysaccharides, nucleic acids, proteins, lipids, and inorganic ions^1^. When left uncontrolled, dental plaque can cause gingivitis, periodontitis, and caries^2,3^. Caries is a major global health problem estimated to affect 3 billion people worldwide^4^ and disproportionately affecting people with lower socioeconomic status^5^. In addition to these oral diseases, poor oral health is also associated to several other systemic diseases^2,6^.

Enzymes that target and break down the extracellular matrix show potential for controlling dental biofilms without disturbing the healthy microbiota and hence promoting oral health^7^. Glucanohydrolases, such as dextranases and mutanases, target the polysaccharide matrix and impact dental biofilms^8–13^. Other enzymes that have been tested include lysozyme, beta-glucanases, lipases, proteases, and DNase I^9,10,12,14–18^. Many of these studies report successful prevention and removal of dental biofilms, especially with combinations of enzymes that target different matrix components simultaneously.

Many studies focus on the removal of polysaccharides for dental biofilm control, but eDNA is also a major dental biofilm component^18–22^ and is thus a potential target for enzyme-based biofilm control. It is well documented that DNase I removes young biofilms but is ineffective against mature biofilms^23^. This has been shown for many types of biofilm^24^, including dental biofilms^14,18,25,26^. The reason for this discrepancy was only recently uncovered and involves the presence of DNA structures that are resistant to DNase I, an endonuclease with preference for cleaving at AT base pairs in dsDNA, and with some (although strongly reduced) activity for cleaving ssDNA and DNA-RNA hybrids.

Extracellular DNA in biofilms originates from genomic DNA from the bacteria or immune cells. It was therefore thought to exist as B-DNA: the canonical right-handed double helix characterised by Watson-Crick base pairing, which is sensitive to DNase I^27^. However, DNA can also adopt noncanonical secondary structures, such as Z-DNA, G-quadruplexes (G4), i-Motifs, and triplex-DNA. Among these structures, Z-DNA and G4 can be abundant in biofilms^28–30^.

Z-DNA is a left-handed double helix that preserves Watson-Crick base pairing and resists degradation by DNase I^31^. It is stabilised in biofilms by DNA-binding proteins belonging to the DNABII family, and Z-DNA has been detected in a variety of biofilms *in vivo* and *in vitro*^28,29,32^. G4 structures are characterised by noncanonical Hoogesteen base pairing in nucleotide regions rich in guanine. G4-DNA and -RNA have been detected *in vitro* in biofilms formed by *Pseudomonas aeruginosa*^30^, *Staphylococcus epidermidis*^28^, and *in vivo* in an implant-associated *Staphylococcus aureus* infection^28^. G4 structures also resist degradation by DNase I^28^.

The discovery of non-canonical DNA structures in biofilms is so recent that only few studies have investigated if oral microorganisms produce such structures in the biofilm matrix. Z-DNA has been found in the matrix of *Streptococcus mutans* biofilms and in dental calculus^29,32^, but it is currently unknown whether G4 and Z-DNA structures exist in *in vivo*-grown dental biofilms. We hypothesised that these structures are present in dental biofilms and that they impede biofilm removal by DNase I. In this study, we investigate the presence of G4 and Z-DNA in dental biofilms from healthy and caries-active subjects, and we thereby assess whether these are relevant targets for improved dental biofilm control.

## Methods

### Materials, bacterial strains, and growth conditions

For fluorescence microscopy experiments, immunolabelling was used to visualise G4, Z-DNA, and B-DNA using a modified version of the protocol from Minero *et al.*^28^. BG4 goat IgG with Atto-488 conjugation (Ab00174-24.1, Absolute Antibody) was used to visualise G4 structures, and Z22 rabbit IgG with Atto-488 conjugation (Ab00783-23.0, Absolute Antibody) was used to visualise Z-DNA. The fluorophore was the same and therefore G4 and Z-DNA were visualised independently. B-DNA was visualised simultaneously with either G4 or Z-DNA using a primary Anti-dsDNA mouse IgG (ab27156, Abcam) paired with a secondary goat anti-mouse IgG with Alexa Fluor 405 Plus conjugation (A48255, Invitrogen). The primary BG4, Z22, and Anti-dsDNA antibodies were diluted 1/100 in blocking buffer (3 % bovine serum albumin (BSA; A7906, Sigma-Aldrich) in 1 x phosphate buffered saline (PBS; E703, VWR)), and the secondary antibody was diluted 1/150 in blocking buffer. Microorganisms were visualised using the membrane stain FM 4-64 (10 µg/ml in 1 x PBS; T13320, Invitrogen). DNase I (04716728001, Roche) was used at 500 U/mL in a buffer for optimal enzyme activity (25 mM Tris-HCl (RES3098T-B7, Sigma-Aldrich), 6.25 mM CaCl_2_ (223506, Sigma-Aldrich), 1 mM MgSO_4_ (M1880, Sigma-Aldrich), pH 7.5)^28^.

### Collection of dental biofilms

Supragingival plaque was collected from 10 healthy volunteers with no clinical signs of periodontal disease or active caries lesions, and from 10 caries-active patients with at least three active caries lesions who were undergoing treatment for caries at the Department of Dentistry and Oral Health, Aarhus University. The Nyvad criteria and the periodontal screening index were used to assess the caries and periodontal status, respectively^33,34^. Samples were collected between August 2023 and May 2024. Pooled plaque was collected from all buccal tooth surfaces of the first quadrant with a universal curette and transferred to a sterile Eppendorf tube. Plaque was used for experiments immediately after collection. All volunteers signed informed consent forms. The protocol was approved by the Ethical Committee of Region Midtjylland (case no. 1-10-72-178-18).

### Visualisation of G4, Z-DNA, and B-DNA in dental biofilms via immunolabelling

To visualise G4, Z-DNA, and B-DNA, confocal laser scanning microscopy (CLSM) of fluorescently conjugated antibodies specific to these DNA structures was used. All immunolabelling was performed on live, fresh dental biofilms that were processed and imaged immediately after collection. The protocol was modified from Minero *et al*. 2024^28^, all steps were performed at room temperature, and all incubations were performed in the dark.

A hydrophobic marker was used to draw a small ring on positively charged microscope slides (Superfrost, 631-0108, VWR). Each dental biofilm sample was split in two, and each half was transferred to separate microscope slides to visualise G4 and Z-DNA in the same sample. Samples were first blocked with 60 μL blocking buffer for 30 min, then hybridised with 60 µL BG4 or Z22 antibody in blocking buffer for 1 h before addition of 60 μL anti-dsDNA antibody and further incubated for 1 h. The samples were washed by removing the liquid and replacing it with 120 μL 1 x PBS twice before addition of 60 μL Alexa Fluor 405 Plus-conjugated secondary antibody and incubation for 1 h to label the anti-dsDNA primary antibody. The liquid was then discarded, and samples were washed twice with 120 µL 1 x PBS. Finally, 30 µL FM 4-64 was added and incubated in the dark for 5 min. A microscope coverslip was placed on top of the samples and the edges were sealed with nail polish. The samples were imaged immediately with CLSM (LSM700, Zeiss) using a Plan-Apochromat 63x/1.40 NA oil immersion objective lens with 5 mW 405 nm and 10 mW 488 nm lasers operating at 3 % and 4.5 % power, respectively. The samples were highly variable, and thus fields of view (FOV) were chosen to show locations of the plaque which contained G4 or Z-DNA, and notes were kept to qualitatively describe the appearance of the biofilms. A minimum of 3 FOV were imaged per sample, which were a mixture of Z-stacks and 2D snapshots. Imaging settings were varied due to variations in fluorescence intensity in different biofilm locations by adjusting the master gain and pixel dwell time. Image brightnesses in the figures were therefore adjusted individually in Fiji ImageJ^35^ for presentation purposes, and the data is descriptive of whether different eDNA structures were present in the samples and is not quantitative.

### DNase I treatment of dental biofilm

Supragingival plaque was collected from one subject with no clinical signs of periodontal disease or active caries lesions whose plaque contained G4 and Z-DNA structures. The sample was distributed into multiple tubes and suspended in 100 µL PBS. The PBS was removed and replaced by either 200 µL DNase I (500 U/mL) diluted in buffer or 200 µL buffer with no enzyme. Samples were incubated for 1 h at 37 °C, followed by immunolabelling to visualise G4, Z-DNA, B-DNA, and microorganisms as described above. Immunolabelling was performed in tubes rather than on microscope slides. Finally, the labelled samples were transferred to microscope slides and visualised by CLSM (LSM700, Zeiss) using a Plan-Apochromat 63x/1.40 NA oil immersion objective lens with 5 mW 405 nm and 10 mW 488 nm lasers operating at 3 % and 4.5 % power, respectively. The imaging settings for visualising G4, Z-DNA, and B-DNA were kept the same when imaging samples with and without DNase I treatment so that the conditions could be compared. At least 5 FOV were acquired per sample and the experiment was performed in triplicate on separate days.

## Results

### G4 and Z-DNA are present in dental biofilms from healthy subjects

We visualised G4, Z-DNA, and B-DNA in dental biofilms from healthy subjects to assess whether noncanonical DNA structures are present in habitual plaque (Figure 1, Figure 2). We detected large areas of eDNA rich in G4 structures surrounding clusters of microorganisms (Figure 1A-E) in dental biofilms from 6 out of 10 subjects, and large eDNA structures containing Z-DNA in 9 out of 10 (Figure 2A-D). There was considerable variation in the abundance of eDNA and non-canonical DNA structures within a given plaque sample and between samples from different individuals. For most subjects, the structures represented a small fraction of the total biofilm, although a few samples appeared to contain large amounts. Quantification was not possible due to the large sample heterogeneity, and we therefore chose to qualitatively assess the appearance of the biofilms.

eDNA containing G4 or Z-DNA appeared to be associated to groups of microorganisms with distinctive morphologies (Figure 1B, 1C, 2B, 2C). In general, these microorganisms were cocci or rods of various sizes. Some clusters were surrounded by concentric rings of eDNA formed by an inner layer of Z-DNA or G4 and an outer layer of B-DNA (Figure 1B, 1D, 2A, 2B). Other clusters were surrounded by a ring of G4, Z-DNA, or B-DNA alone and not in combination with other eDNA structures (Figure 1D, 2C, 2D). We are critical as to whether the antibodies used to visualise the different structures were able to fully penetrate the biofilm; thus, fluorescence around the rim of bacterial clusters could indicate that antibodies only bound the outer layers of the biofilm matrix. However, since B-DNA was always the outermost layer, it would imply that the antibody used to visualise B-DNA penetrated biofilms less than other antibodies, which is unlikely.

In addition to the large eDNA structures in bacterial clusters, we also observed individual microorganisms with smaller quantities of G4 or Z-DNA bound to the cell surface. These were present in plaque from 7 out of 10 subjects for G4, and in 9 out of 10 subjects for Z-DNA. In general, individual microorganisms with surface-associated secondary DNA structures were either small rods or were very large and long cells that resembled fungal hyphae (Figure 1F, 2E, 2F).

**Figure 1.**
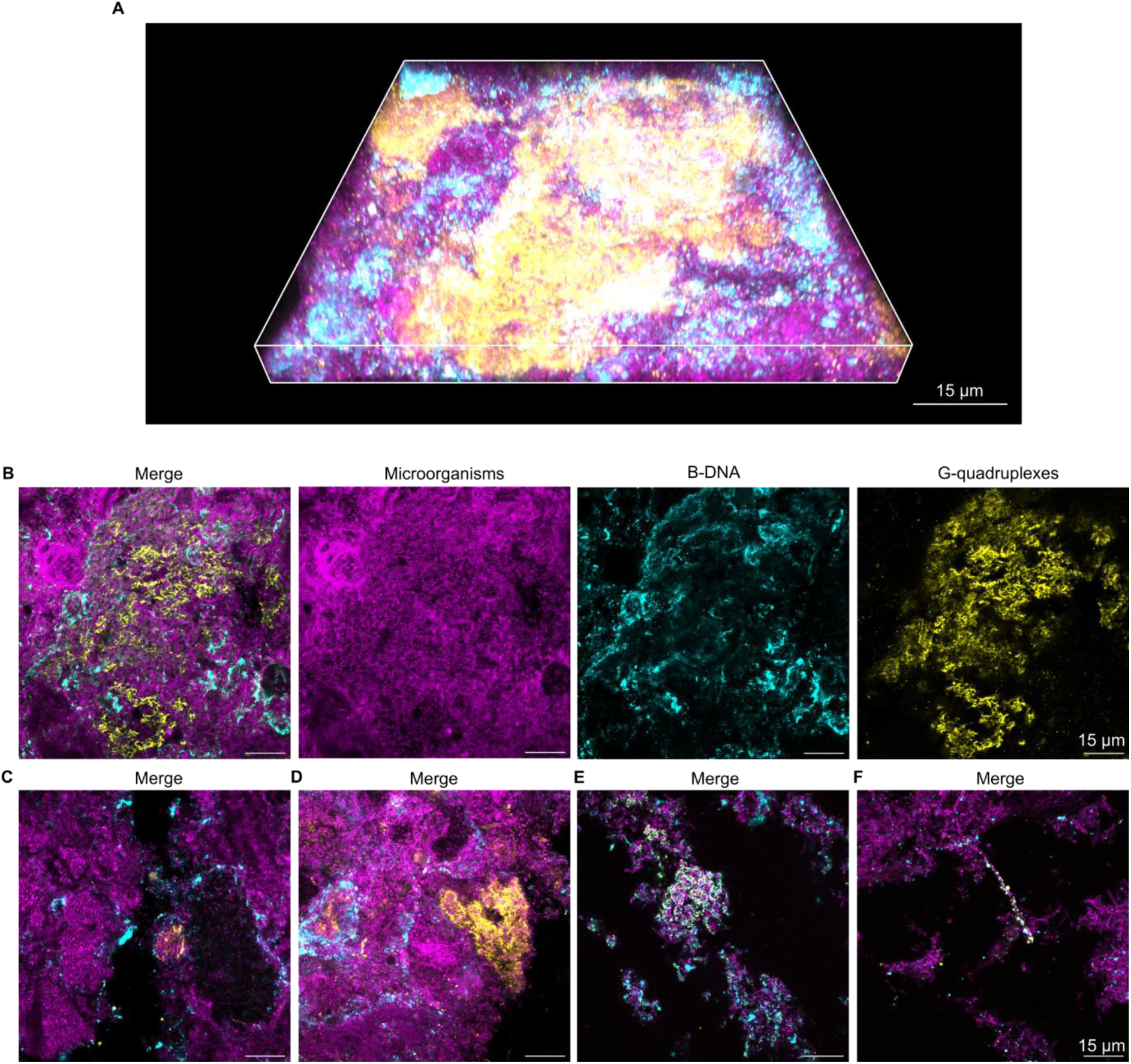
Extracellular G4 structures in dental biofilms from healthy subjects. Confocal microscopy images show microorganisms (magenta), B-DNA (cyan), and G4 (yellow) in dental biofilms. **A)** 3D merged confocal image of an example of a G4 structure in a dental biofilm from a healthy subject. **B)** Single channel and merged 2D confocal images of G4 structures in a dental biofilm from a healthy subject. **C-F)** Merged 2D images showing further examples of G4 in dental biofilms from healthy subjects. The image in A) was prepared in Zen Blue (Zeiss) and the images in C-F) are images of single planes in biofilms that were prepared in Fiji ImageJ^35^.

**Figure 2.**
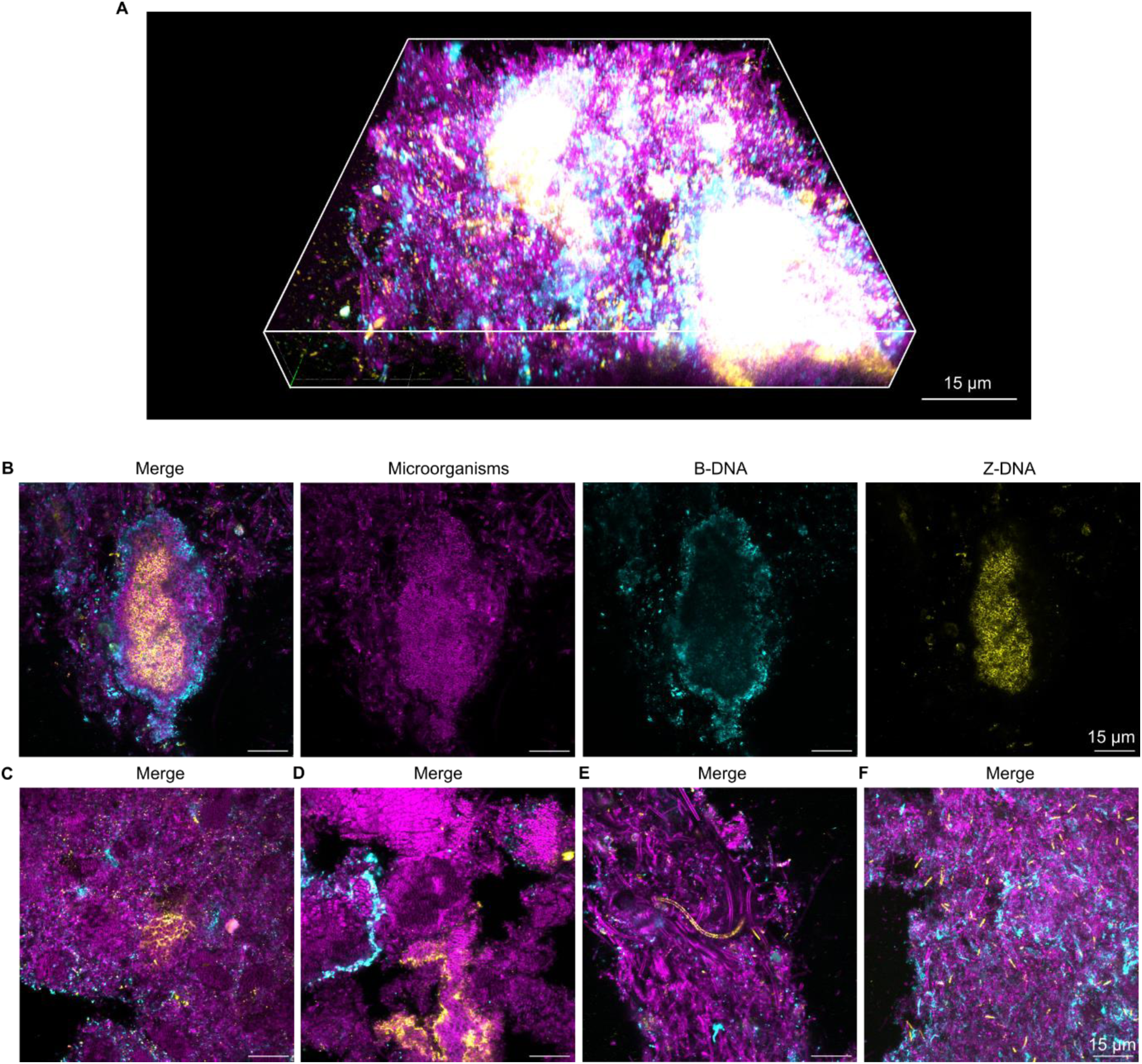
Extracellular Z-DNA structures in dental biofilms from healthy subjects. Confocal microscopy images show microorganisms (magenta), B-DNA (cyan), and Z-DNA (yellow) in dental biofilms. **A)** 3D merged confocal image of an example of a Z-DNA structure in a dental biofilm from a healthy subject. **B)** Single channel and merged 2D confocal images of a Z-DNA structure in a dental biofilm from a healthy subject. **C-F)** Merged 2D images showing further examples of G4 in dental biofilms from healthy subjects. The image in A) was prepared in Zen Blue (Zeiss) and the images in C-F) were prepared in Fiji ImageJ^35^. C-D) are images of single planes within biofilms, and F) is a maximum intensity Z-projection.

### G4 and Z-DNA are present in dental biofilms from caries-active subjects

Dental biofilms from healthy subjects contained both G4 and Z-DNA. Dental biofilms from healthy and caries-active subjects have different microbial compositions, and we therefore investigated if G4 and Z-DNA are also associated with dental biofilms from caries-active subjects. We collected dental biofilms from 10 patients with caries, and visualised G4, Z-DNA, and B-DNA via immunolabelling and CLSM (Figure 3, Figure 4).

We detected large eDNA structures containing G4 (Figure 3A-E) and Z-DNA (Figure 4A-E) surrounding large clusters of microorganisms. Biofilms from 6 out of 10 caries-active subjects contained large eDNA structures containing G4, and biofilms from 7 out of 10 samples contained large Z-DNA structures. These large eDNA structures were associated to groups of microorganisms with different morphologies, including rods and cocci of different sizes (Figure 3B, 3C, 4B, 4C). Similar to the dental biofilms from healthy subjects, some bacterial clusters contained eDNA in concentric rings with an outer layer of B-DNA and an inner layer of G4 or Z-DNA (Figure 3A, 3E, 4B, 4D, 4E), while other clusters contained a more homogenous distribution of eDNA (Figure 3B, 4C). Like in healthy subjects, samples where heterogenous and the large eDNA structures presented in Figures 3 and 4 only represent a small fraction of the biofilm.

We also detected individual microorganisms with surface-associated G4 or Z-DNA structures: 8 out of 10 dental biofilm samples from caries-active subjects contained individual microorganisms with surface-associated G4 structures, and 6 out of 10 contained microorganisms with surface-associated Z-DNA structures. Generally, these microorganisms resembled fungal hyphae (Figure 3F, 4F).

**Figure 3.**
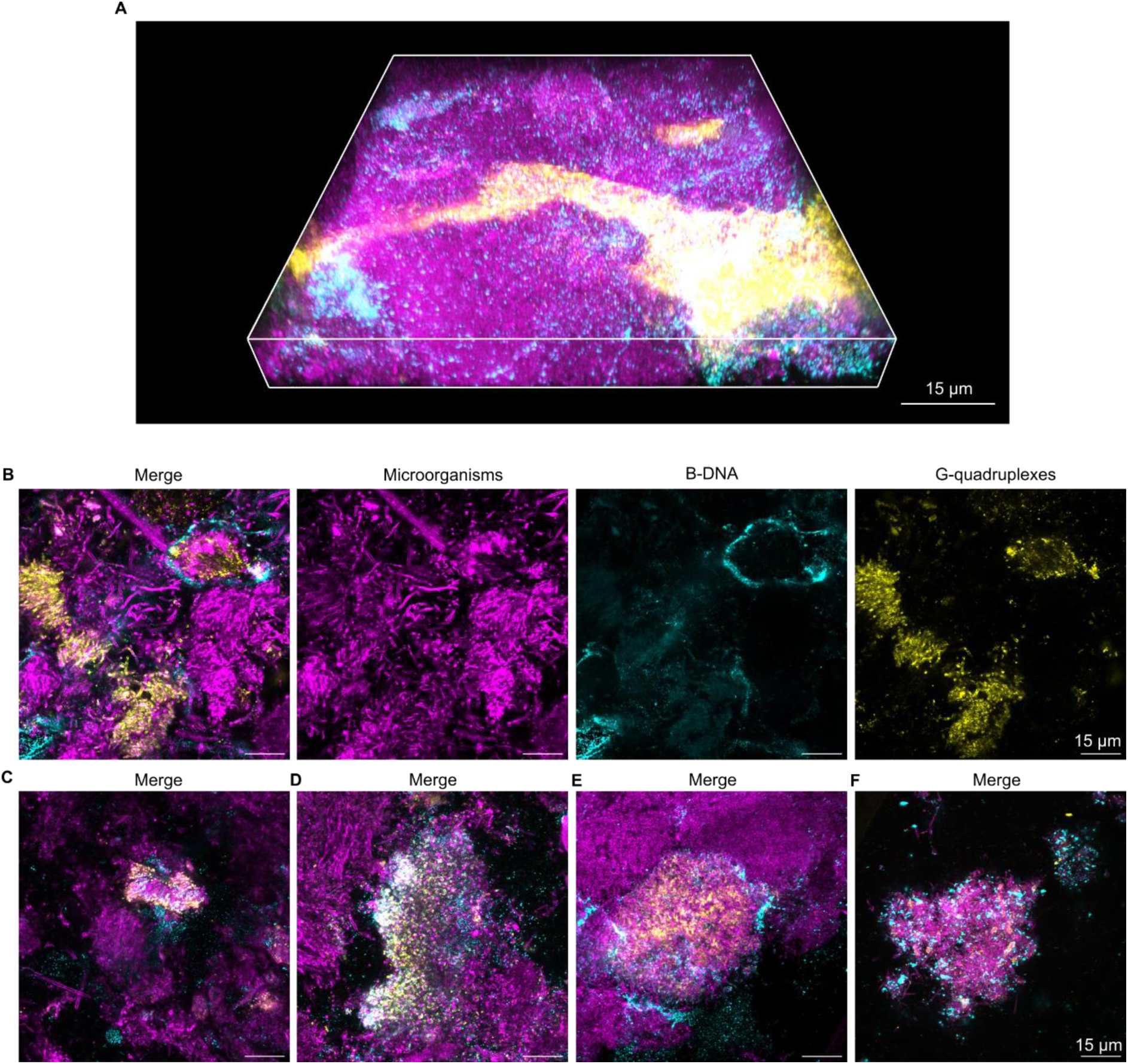
Extracellular G4 structures in dental biofilms from caries-active patients. Confocal microscopy images of microorganisms (magenta), B-DNA (cyan), and G4 (yellow) in dental biofilms from caries-active subjects. **A)** Merged 3D image of an example G4 structure in a dental biofilm from a caries-active subject. **B)** Single channel and merged 2D images of G4 structures in a dental biofilm from a caries-active subject. **C-F)** Merged 2D images of G4 structures in dental biofilms from caries-active subjects. The image in A) was prepared in Zen Blue (Zeiss) and the images in C-F) are images of single planes in biofilms that were prepared in Fiji ImageJ^35^.

**Figure 4.**
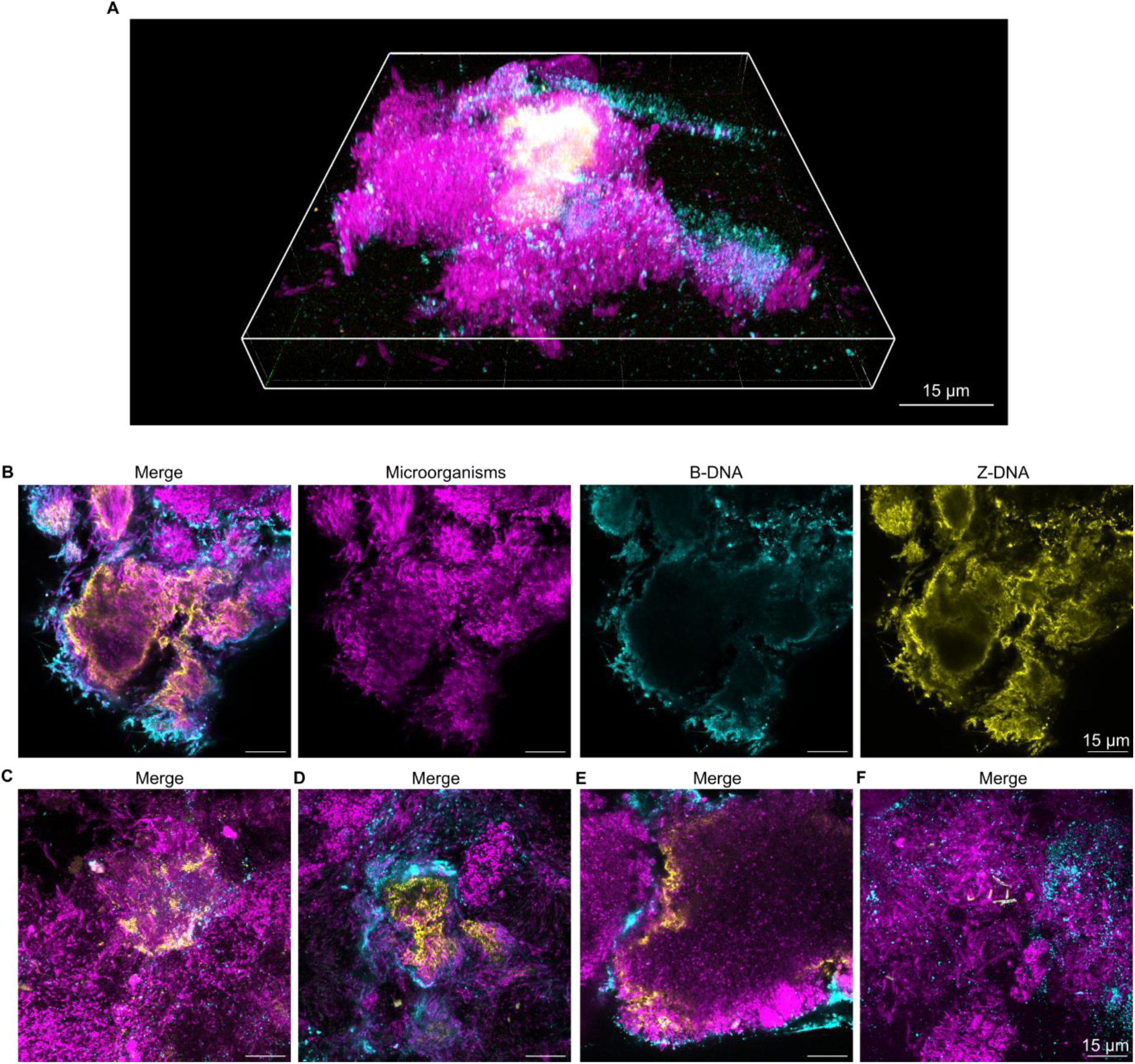
Extracellular Z-DNA structures in dental biofilms from caries-active patients. Confocal microscopy images of microorganisms (red), B-DNA (cyan), and Z-DNA (yellow) in dental biofilms from caries-active subjects. **A)** Merged 3D image of an example Z-DNA structure in a dental biofilm from a caries-active subject. **B)** Single channel and merged 2D images of a Z-DNA structure in a dental biofilm from a caries-active subject. **C-F)** Merged 2D images of Z-DNA structures in dental biofilms from caries-active subjects. The image in A) was prepared in Zen Blue (Zeiss) and the images in C-F) are images of single planes in biofilms that were prepared in Fiji ImageJ^35^.

### G4 and Z-DNA structures remain in dental biofilms after treatment with DNase I

DNase I is often used to degrade eDNA in biofilms, but it is ineffective at removing mature biofilms^23^, possibly due to the resistance of G4 and Z-DNA to degradation by DNase I^28,29^. We investigated whether DNaseI could degrade non-canonical DNA structures in dental biofilms by comparing G4, Z-DNA, and B-DNA in dental biofilms from the same subject after treatment with DNase I or buffer. The subject belonged to the group of 10 participants without clinical signs of caries or periodontitis. We could not compare the same FOV before and after treatment because the antibodies used for immunolabelling would protect the DNA from degradation by DNase I. We therefore compared the presence of G4 and Z-DNA in arbitrarily chosen FOV in separate samples (taken from the same subject) with and without DNase I treatment.

Both G4 and Z-DNA structures were present in the dental biofilm regardless of whether the sample was treated with DNase I (Figure 5). This indicates that G4 and Z-DNA are unaffected by treatment with DNase I. It was expected that DNase I would remove B-DNA in the biofilms, but interestingly, signal from B-DNA appeared very similar in the samples with and without DNase I treatment and was located in regions where it overlapped with either G4 or Z-DNA structures. Perhaps B-DNA was also protected from degradation by these secondary structures or by another matrix component. We confirmed that DNase I was active against DNA in a separate experiment (Supplementary Figure S1), therefore, this is the first indication that DNase I fails to remove B-DNA from biofilms.

**Figure 5.**
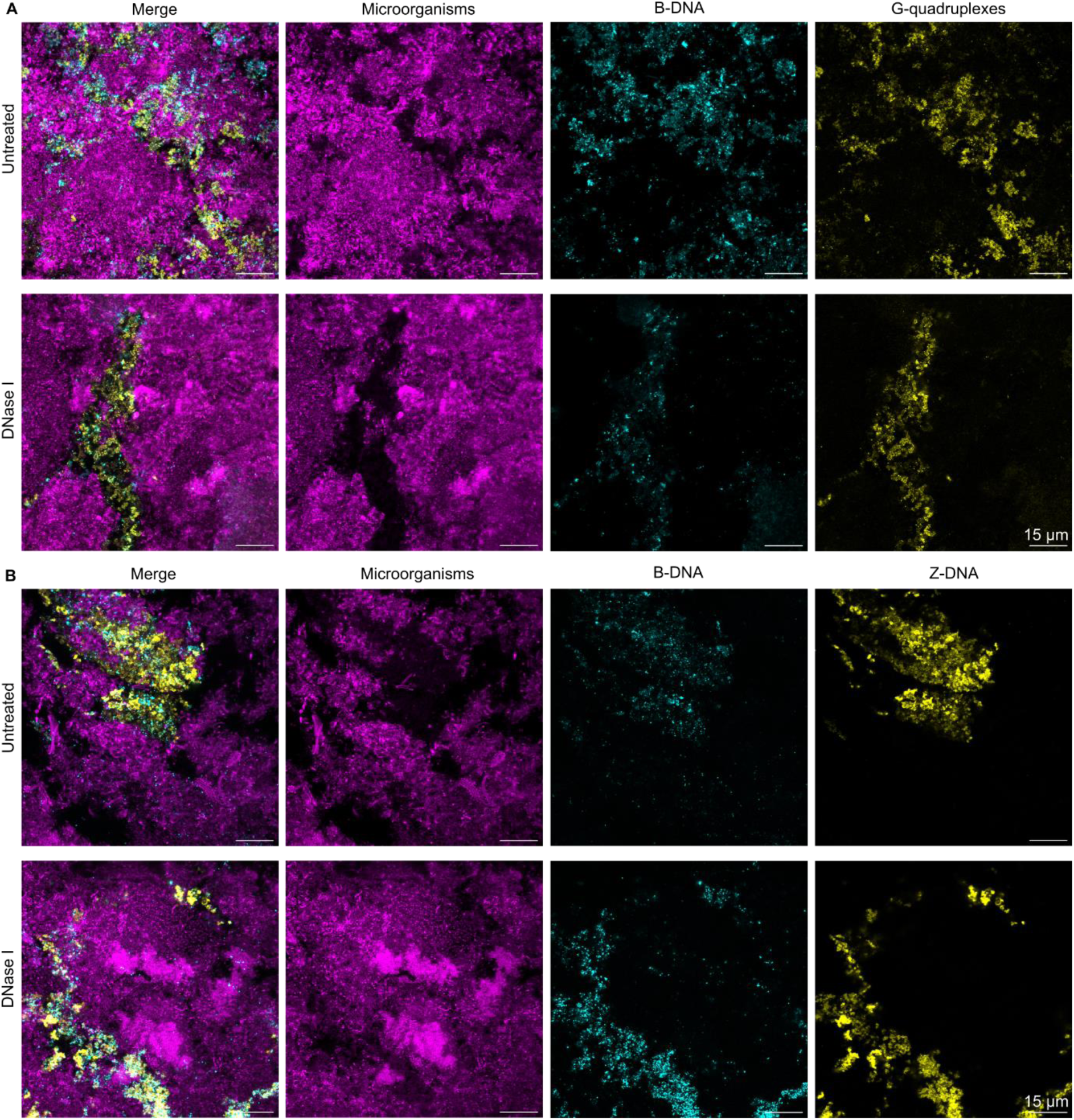
Extracellular DNA structures remain in dental biofilms after treatment with DNase. **I.** Confocal microscopy images of microorganisms (magenta), B-DNA (cyan), and G4/Z-DNA (yellow) in dental biofilms from a healthy subject visualised by confocal microscopy. **A)** Single channel and merged 2D images of G4 in dental biofilms with and without treatment with DNase I. **B)** Single channel and merged 2D images of Z-DNA in dental biofilms with and without treatment with DNase I. G4 and Z-DNA structures remained in the dental biofilms after a 1 h treatment with DNase I. All images in this figure were collected using the same microscope settings, and the brightnesses of all images were adjusted equally in Fiji ImageJ^35^.

## Discussion

eDNA is an important component of the extracellular matrix of dental biofilms^1,18,19^. Our research supports this and furthermore demonstrates that a fraction of eDNA in dental biofilms is in the Z-DNA conformation. For the first time, we show that dental biofilms also contain G4 structures (Figures 1-4). In general, these structures comprised a relatively small fraction of the total dental biofilm in both healthy and caries-active subjects, but they were present in the majority of samples tested from both groups. Secondary eDNA structures filled the extracellular space in clusters of microorganisms (Figure 1B, 1C, 2B, 2C, 3B, 3C, 4B, 4C), and were sometimes also associated to the surfaces of individual cells (Figure 1F, 2E, 2F, 3F, 4F). This indicates that local conditions promote the formation of secondary DNA structures, and that particular species within the diverse microbial community of dental biofilms have the ability to form and associate to such DNA structures.

eDNA in dental biofilms originates both from bacterial cells and from the host^36^, and previous research shows that bacteria manipulate both bacterial and host eDNA to adopt the Z-DNA conformation^29,32^ . Local hot-spots with high concentrations of eDNA in general, and Z-DNA in particular, therefore depend on a number of potential factors. Bacteria acquire eDNA from neighbouring cells through processes of autolysis or allolysis, and eDNA from host cells arrives when neutrophils expel their chromatin as neutrophil extracellular traps (NETs). The conversion of eDNA from the B-to Z-DNA conformation then requires additional local factors, such as DNABII proteins released from bacterial cells. Our analysis was purely qualitative, but further analysis into the local species composition, matrix composition, and presence of e.g. histones associated with NETs could point to the origin of Z-DNA, the local conditions, and the microorganisms responsible for hotspots of Z-DNA and G4 structures in the biofilm. Future in-depth investigations of this kind will no doubt improve our understanding of how these DNase I-resistant structures form in dental biofilms.

The antibody used for detecting G4 in our study binds to both DNA and RNA G4, and likewise, the antibody used to detect Z-DNA also binds Z-RNA. Extracellular RNA has recently been shown to be an important component of some biofilms^37,38^ and therefore it is possible that dental biofilms contain both G4 DNA and RNA, and both Z-DNA and Z-RNA. We expected that G4 DNA would only form in regions that colocalise with B-DNA, yet in some cases G4 were detected in regions that lacked B-DNA (Figure 1D, 3B), which may suggest that some of the G4 structures were comprised of RNA and not DNA.

G4 and Z-DNA structures resist degradation by mammalian DNase I, which has been shown in a number of biofilms, including Z-DNA in dental calculus^28,29,32^. Our results confirm the trend, showing that G4 and Z-DNA remain in dental biofilms after treatment with DNase I (Figure 5). To our surprise, B-DNA associated with these structures also remained in the biofilm after DNase treatment, indicating that DNase I cannot degrade some of the B-DNA within DNA superstructures that contain non-canonical DNA.

Our results highlight that G4 and Z-DNA structures are present in dental biofilms from the majority of subjects, and we find no indication that healthy and caries-active subjects differ in this respect. We collected plaque from the buccal surface, which is easily reached during toothbrushing. We therefore expect that the dental biofilms were relatively young (6-12 hours), and the abundance of eDNA and secondary structures may change if biofilms are left undisturbed for longer periods of time. For example, Buzzo *et al*.^29^ showed in pure culture biofilms that Z-DNA increased in abundance the longer biofilms were left to grow. It is therefore noteworthy that they are present in the majority of dental biofilms sampled in this study.

The high concentration of Z-DNA and G4 structures in large bacterial clusters shows that non-canonical DNA structures can dominate the matrix composition locally. This is important to notice because even local patches of highly resilient biofilm can affect oral health. Wen *et al*.^32^ showed how the resilience of Z-DNA led to long-term build-up of eDNA, causing calculus formation. Our discovery of Z-DNA and G4 hotspots in dental biofilms raises new questions of how these structures affect biofilm resilience and oral health. It is not always the biofilm at large, but rather sub-structures with a specific phenotype that cause oral disease. Kim *et al*.^39^ showed that bacterial clusters of *S. mutans* surrounded by a rim of *Streptococcus oralis* were responsible for acid production and enamel dissolution leading to caries. The two populations were distinctly segregated in a core-shell structure connected via a scaffold of extracellular matrix produced by *S. mutans. S. mutans* does produce eDNA in the biofilm matrix, and *in vitro* analyses reveal Z-DNA in pure culture biofilms of the species^29^.

Having observed several bacterial clusters with a core-shell structure of Z-DNA/B-DNA in dental biofilms, we propose to pursue investigations of how non-canonical DNA structures might contribute to more virulent sub-phenotypes within dental biofilms.

## Conclusion

We have demonstrated for the first time the presence of G4 and Z-DNA in the extracellular matrix of dental biofilms from both healthy and caries-active subjects. The majority of samples from both groups contained large eDNA structures containing G4 or Z-DNA, as well as individual microorganisms with surface-associated G4 or Z-DNA. G4 and Z-DNA structures remained in dental biofilms after treatment with DNase I. G4 and Z-DNA structures are potentially novel targets for developing improved enzyme-based products for oral care.

## Author contribution statement

RLM conceptualised the work. RLM, SS, and MKT acquired funding. RLM, SS, LGP, IK and MKT supervised the work. DCSE designed and performed experiments and drafted the original manuscript. MFK performed experiments and collected patient samples. All authors discussed the results, and all authors contributed to the final manuscript.

## Data availability statement

The datasets used during the current study are available from the corresponding author on reasonable request.

## Acknowledgements

This research was funded by a grant (2051-00007B) from the Innovation Foundation Denmark in partnership with Novonesis A/S Denmark. We would like to say a big thank you to Pernille Rikvold for her valuable contribution to plaque collection.

## Verifying DNase I activity

To verify DNase I activity, we treated salmon sperm DNA with DNase I and compared it to an untreated sample by visualising the DNA on an agarose gel. Salmon sperm DNA (100 μg, D1626, Sigma-Aldrich) was incubated for 2 h at 37 °C with DNase I (500 U/mL) in buffer in a reaction volume of 100 µL. A control sample of salmon sperm DNA (100 μg) was incubated under the same conditions in 100 µL buffer with no enzyme. 100 ng each of the DNase I-treated and control sample were loaded onto an agarose gel (1 % agarose in 0.5 X TBE buffer) alongside a ladder (TrackIt 1 kb Plus DNA Ladder, 10488085, Invitrogen), and gel electrophoresis was performed at 120 V for 1.5 h. DNA in the gel was stained by Sybr SAFE DNA Gel Stain (S33102, Invitrogen) and visualised in a Gel Doc EZ Imager (BioRad) with Image Lab Software (Biorad).

A band containing DNA was only seen in the lane that was loaded with untreated DNA, and not in the lane that was loaded with the DNase I-treated DNA (Figure S1). This verifies that the DNase I was active against DNA.

**Figure S1.**
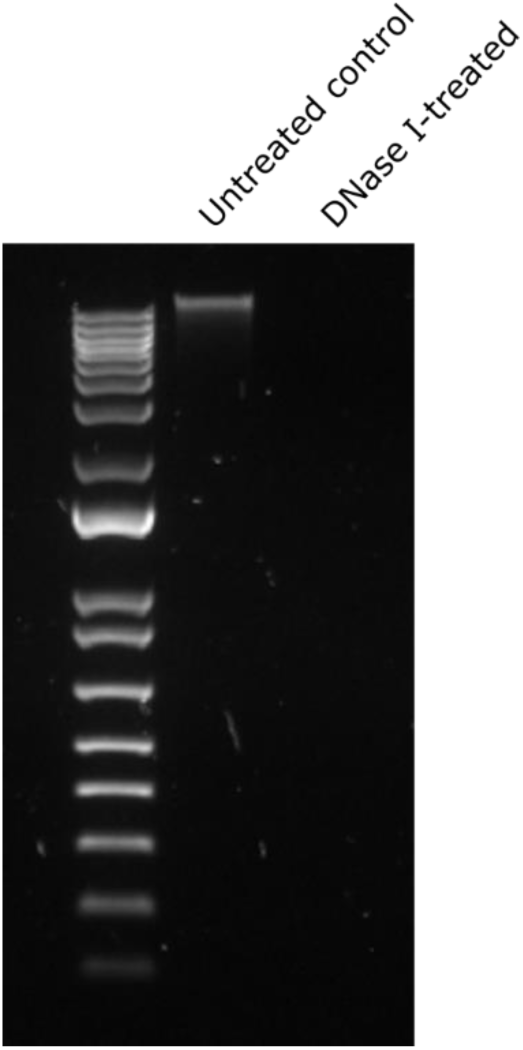
Salmon sperm DNA either treated with DNase I (right lane) or untreated (left lane). DNA was treated in buffer for a total time of 2 h at 37 °C. All DNA was removed by DNase I during this incubation.

